# Setd2 inactivation sensitizes lung adenocarcinoma to inhibitors of oxidative respiration and mTORC1 signaling

**DOI:** 10.1101/2022.10.06.511146

**Authors:** David M. Walter, Amy C. Gladstein, Katherine R. Doerig, Ramakrishnan Natesan, Saravana G. Baskaran, A. Andrea Gudiel, Keren M. Adler, Jonuelle O. Acosta, Douglas C. Wallace, Irfan A. Asangani, David M. Feldser

## Abstract

*SETD2* is a tumor suppressor that is frequently inactivated in several cancer types. The mechanisms through which SETD2 inactivation promotes cancer are unclear, and whether targetable vulnerabilities exist in these tumors is unknown. Here we identify heightened mTORC1-associated gene expression programs and functionally higher levels of oxidative metabolism and protein synthesis as prominent consequences of *Setd2* inactivation in KRAS-driven mouse models of lung adenocarcinoma. Blocking oxidative respiration and mTORC1 signaling abrogates the high rates of tumor cell proliferation and tumor growth specifically in SETD2-deficient tumors. Our data nominate SETD2 deficiency as a functional marker of sensitivity to clinically actionable therapeutics targeting oxidative respiration and mTORC1 signaling.

## Introduction

Inactivation of *SETD2* is a prevalent feature of many cancer types including lung adenocarcinoma [1–3]. We and others have shown that inactivation of *Setd2* in *Kras^G12D^*-driven mouse models of lung adenocarcinoma promotes rampant cellular proliferation very early after tumor initiation; an effect that is widespread amongst all developing tumors [4, 5]. Moreover, *Setd2* quantitatively ranks at the apex of all major tumor suppressors for its ability to suppress KRAS-driven cell proliferation [4, 6]. However, unlike other tumor suppressors such as *Trp53* or *Rb1*, whose inactivation promotes cell state changes that drive malignant progression, loss of differentiation, and metastasis, inactivation of *Setd2* seems only to fuel cellular proliferation in these models [14–16]. As such, we sought to interrogate the consequences of *Setd2* inactivation in KRAS-driven lung adenocarcinoma in order to better understand how SETD2 deficiency drives early and widespread tumor growth.

## Results

To gain mechanistic insights into how SETD2 constrains cancer growth, we performed a comprehensive analysis of RNA-sequencing data across multiple human cancer types expressing low or high levels of *SETD2* [2]. Using GSEA network analysis, we identified a preponderance of ribosomal- and mitochondrial-associated gene sets that negatively correlated with *SETD2* expression. This association was strongly apparent in multiple tumor types that feature a high frequency of SETD2 mutations, including lung adenocarcinoma (**Fig. 1A and Supplementary Fig. S1A-D**)[7]. The negative correlation of ribosomal- and mitochondrial-associated gene sets was specific to SETD2 expression, as these genes sets were not associated with expression levels of several other major tumor suppressors in lung adenocarcinoma such as *TP53* or *RB1*, which instead correlated with DNA replication-associated gene sets (**Supplementary Fig. S1E-G**). These results suggest that ribosomal- and mitochondrial-biosynthetic pathways are activated in SETD2-deficient tumors and may be responsible for driving rampant tumor cell proliferation. To extend this analysis, we analyzed gene expression data from stage-matched *K-Ctrl* or *K-Setd2^KO^* tumors as well as more advanced, stage-matched tumors isolated from *Kras^LSL-G12D/+^; p53^flox/flox^ (KP)* mice initiated with a nontargeting control lentiviral CRISPR vector (*KP-Ctrl*) or an sgRNA targeting *Setd2 (KP-Setd2^KO^*). Both *K-Setd2^KO^* tumors, which were low grade adenomas, and *KP-Setd2^KO^* tumors, which were higher grade adenocarcinomas, had a strong enrichment of ribosomal- and mitochondrial-associated gene sets over their *K-Ctrl* and *KP-Ctrl* counterparts (**Fig. 1B,C**). Collectively these results suggest that ribosomal- and mitochondrial-biosynthetic pathways are activated in SETD2-deficient tumors and may be responsible for driving rampant tumor cell proliferation.

**Figure 1:**
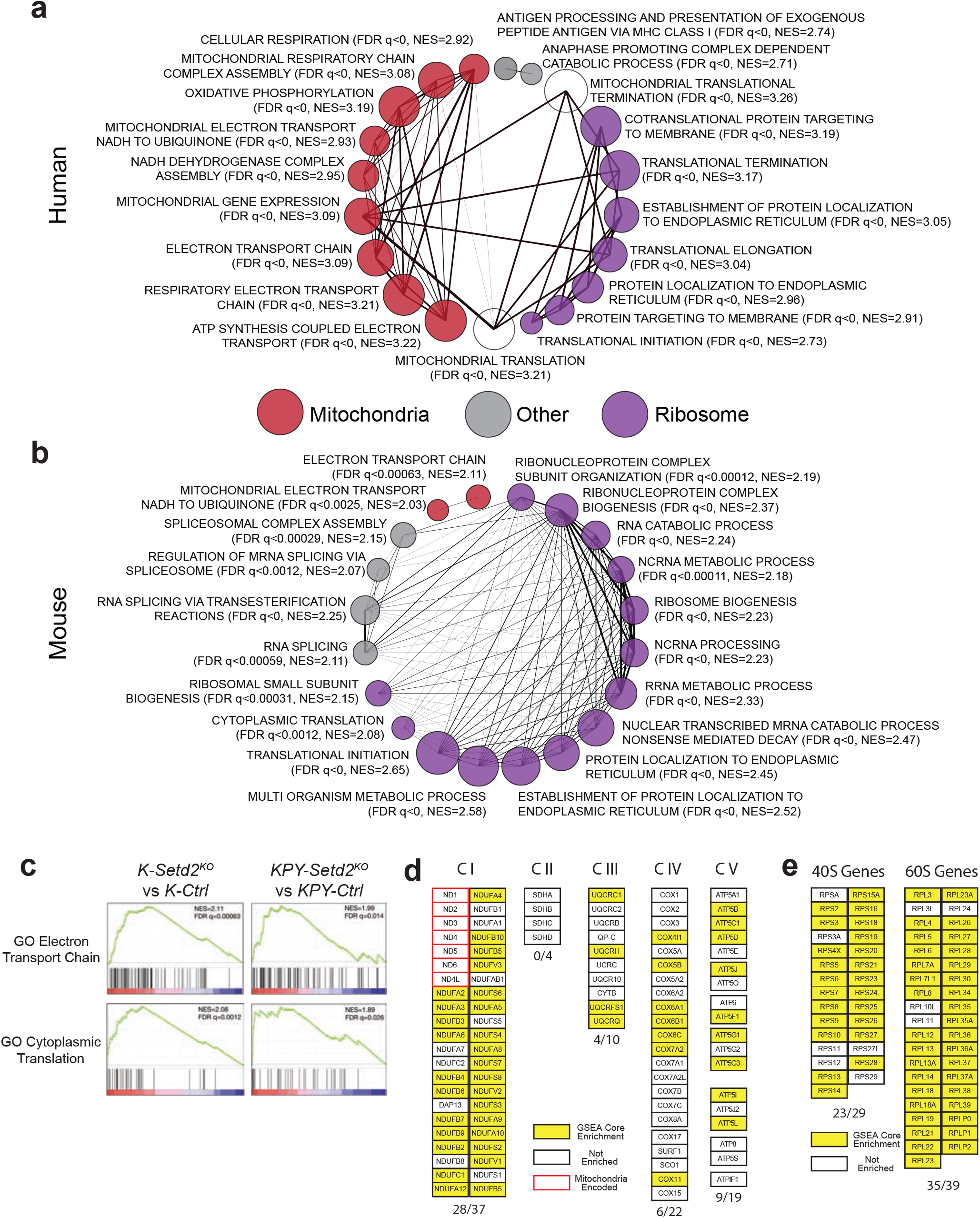
SETD2 deficiency promotes OXPHOS and protein synthesis gene expression programs in human and murine lung adenocarcinomas. **a)** Gene set enrichment analysis (GSEA) network plot of the 20 most enriched gene sets negatively-correlated with SETD2 expression in human lung adenocarcinomas. The size of each node corresponds to the normalized enrichment score (NES) and the width of the connecting lines indicates the number of overlapping genes between gene sets. Gene sets are categorized as mitochondrial, ribosomal or other according to the genes represented. **b)** GSEA network plot of the top 20 enriched gene sets in *K-Setd2^KO^* lung adenocarcinomas. The size of each node corresponds to the normalized enrichment score (NES) and the width of the connecting lines indicates the number of overlapping genes between gene sets. Gene sets are categorized as mitochondrial, ribosomal or other according to the genes represented. **c)** Representative mitochondrial (GO Electron Transport Chain) and ribosomal (GO Cytoplasmic Translation) GO biological process ontology gene sets that are enriched in *K-Setd2^KO^* (left) and *KPY-Setd2^KO^* (right) tumors. **d)** Depiction of mitochondrial electron transport chain genes that show enrichment in the molecular signatures database (mSigDB) Mootha VOXPHOS gene set in *K-Setd2^KO^* tumors [29]. Genes are grouped according to the relevant ETC complex. Genes that are part of the core enrichment of the gene set are marked in yellow, genes that are not enriched are marked in white, and mitochondrial encoded genes are outlined in red. **e)** Depiction of ribosomal 40S and 60S genes that show enrichment in the mSigDB GO Ribosome Biogenesis gene set in *K-Setd2^KO^* tumors. Genes that are enriched in this gene set are marked in yellow while genes that are not enriched are marked in white.

The major driver of enrichment for mitochondrial-associated gene sets in SETD2-deficient tumors was the significant up-regulation of genes encoding mitochondrial electron transport chain (ETC) proteins in complexes I, III, IV and V (**Fig. 1D**). To determine whether physical changes occur in the mitochondria when *Setd2* is inactivated, we imaged *KP-Ctrl* and *KP-Setd2^KO^* tumors by transmission electron microscopy. Although the overall frequency of mitochondria observed per field of view was similar between genotypes, the mitochondria in *KP-Setd2^KO^* tumors were significantly different than those found in *KP-Ctrl* tumors by several parameters (**Supplementary Fig. 2A,B**). Grossly, the mitochondria of *KP-Setd2^KO^* tumors were significantly smaller and had a more electron dense matrix than controls (**Fig. 2A-C Supplementary Fig. 2A**). Morphologically, *KP-Setd2^KO^* mitochondria also had significantly more cristae overall, and on average each cristae was significantly more swollen (**Fig. 2A,D,E**) [8, 9]. While some of these morphological changes can be associated with mitochondrial dysfunction and cell death, the higher electron density in the mitochondria matrix and a greater overall number of cristae in the mitochondria of *KP-Setd2^KO^* tumors may suggest greater oxidative function [10].

**Figure 2:**
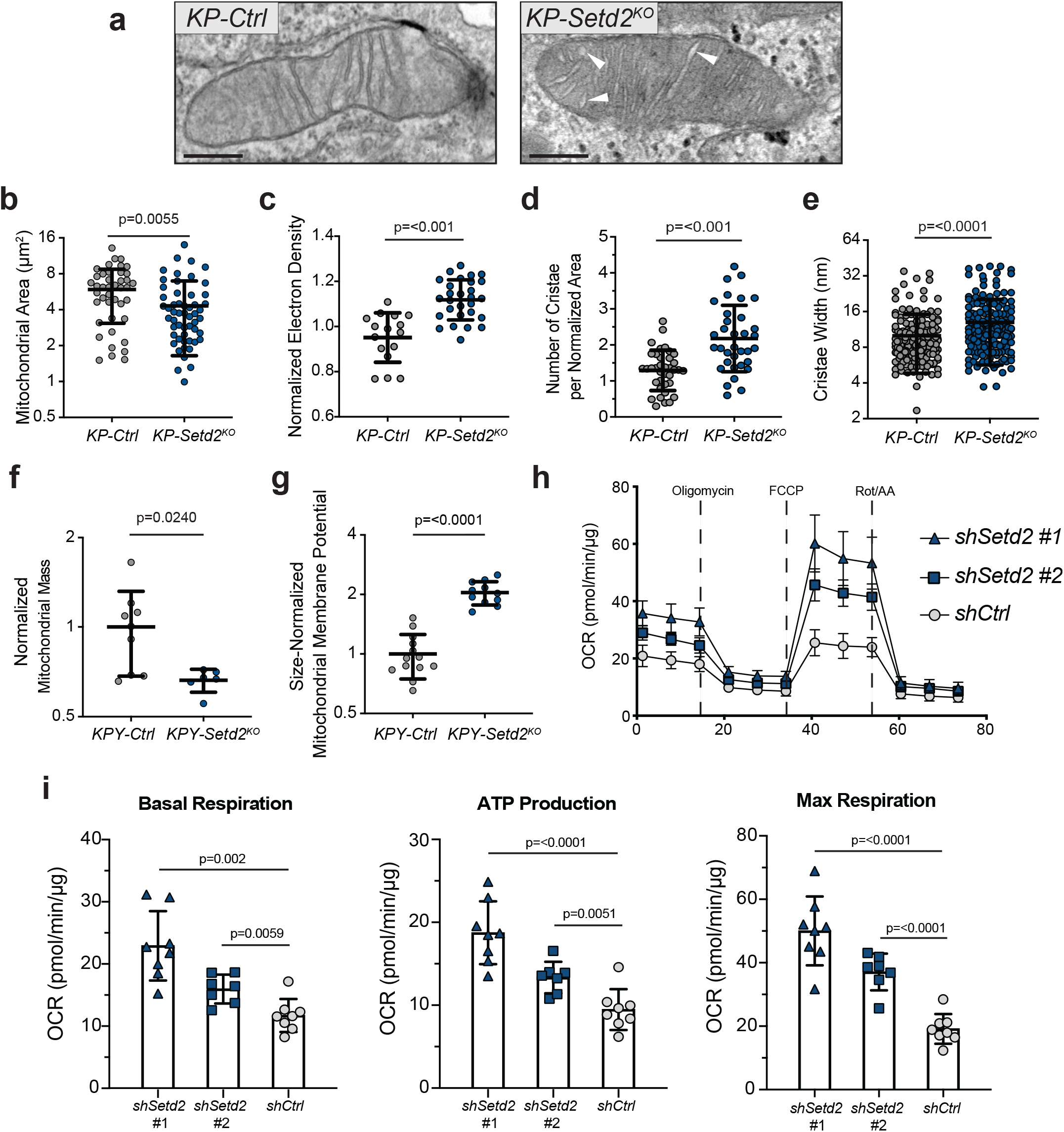
SETD2-deficient tumors contain mitochondria with distinct morphological features and increased electron transport chain activity. **a)** Transmission electron micrograph of mitochondria from a *KP-Ctrl* tumor (left) or *KP-Setd2^KO^* tumor (right). White arrows denote swollen mitochondrial cristae. Scale bars= 250 nm. **b)** Quantification of the area of individual mitochondria measured from electron micrographs. Data indicate the mean ± standard deviation. Data points represent individual mitochondria (*KP-Ctrl*: n=42 mitochondria, n=3 mice, *KP-Setd2^KO^*: n=54 mitochondria, n=3 mice). Significance determined by unpaired Student’s *t*-test. **c)** Quantification of the matrix electron density of individual mitochondria measured from electron micrographs. Data indicate the mean ± standard deviation. Data points represent individual mitochondria (*KP-Ctrl*: n=16 mitochondria, n=2 mice, *KP-Setd2^KO^*: n=28 mitochondria, n=2 mice). Significance determined by unpaired Student’s *t*-test. **d)** Quantification of the number of cristae per mitochondria normalized to mitochondrial area. Data indicate the mean ± standard deviation. Data points represent individual mitochondria (*KP-Ctrl*: n=37 mitochondria, n=3 mice, *KP-Setd2^KO^*: n=34 mitochondria, n=3 mice). Significance determined by unpaired Student’s *t*-test. **e)** Quantification of the width of individual mitochondrial cristae measured from electron micrographs. Data indicate the mean ± standard deviation. Data points represent individual cristae (*KP-Ctrl*: n=179 cristae, n=42 mitochondria, n=3 mice, *KP-Setd2^KO^*: n=177 cristae, n=50 mitochondria, n=3 mice). Significance determined by unpaired Student’s *t*-test. **f)** Quantification of the total mitochondrial mass by median fluorescence intensity of MitoTracker Deep Red FM within tumor cells isolated from *KPY* mice. Data represent the mean ± standard deviation. Data points represent individual tumors (*KP-Ctrl*: n=9 tumors, n=2 mice, *KP-Setd2^KO^*: n=6 tumors, n=2 mice). Significance determined by unpaired Student’s *t*-test. **g)** Quantification of the mitochondrial membrane potential by median fluorescence intensity of MitoProbe DiIC1(5) within tumor cells isolated from *KPY* mice. Data is normalized to the mitochondrial mass of each sample. Data indicate the mean ± standard deviation. Data points represent individual tumors (*KP-Ctrl*: n=13 tumors, n=2 mice, *KP-Setd2^KO^*: n=11 tumors, n=2 mice). Significance determined by unpaired Student’s *t*-test. **h)** Seahorse XF cell mitochondrial stress test assay performed in H2009 sh*Setd2* and sh*Ctrl* cells. Relative oxygen consumption rate was normalized to total protein abundance. Each symbol in OCR profile plots represents the mean of at least n=6 technical replicates of three reading cycles. **i)** OCR profile plots from (**h**) for basal respiration, ATP production, and maximal respiration. Each symbol represents one technical replicate per cell line. Significance determined by unpaired Student’s *t*-test.

To investigate mitochondrial properties in neoplastic cells directly *ex vivo*, we crossed a *Rosa26^LSL-YFP^* Cre-reporter allele into the *KP* model, generating *KPY* mice (**Supplementary Fig. 2C**) [11]. Consistent with the EM data, *KPY-Setd2^KO^* tumor cells had decreased staining for MitoTracker Deep Red indicative of decreased mitochondrial mass (**Fig. 2F**). Though *KPY-Setd2^KO^* tumor cells did not differ from *KPY-Ctrl* tumor cells with respect to mitochondrial superoxide production, the mitochondrial oxidative potential and the total ETC activity was significantly higher in *KPY-Setd2^KO^* tumor cells (**Fig. 2G, Supplementary Fig. 2D,E**).

To determine if the observed mitochondrial changes in SETD2-deficient tumors result in increased mitochondrial function and consequential ATP production, we generated human lung adenocarcinoma cell lines (H2009) with two distinct shRNAs targeting *Setd2. Setd2* shRNA-expressing cell lines had decreased H3K36me3 compared to control cell lines confirming a knockdown of SETD2 (**Supplementary Fig. 3A,B**). To profile mitochondrial function we performed a cell mitochondrial stress test assay (Seahorse XF) on these cell lines. In agreement with our *in vivo* data, SETD2-deficient human cell lines displayed increased oxygen consumption rates and ATP production (**Fig. 2H,I**). These data demonstrate that SETD2-deficiency promotes increased mitochondrial metabolism and are consistent with the gene expression programs that are enriched in human and mouse tumors with low SETD2 expression.

An additional feature of SETD2-deficient tumors was the significant enrichment of genes associated with protein synthesis and ribosome biogenesis (**Fig. 1A-C,E**). To assess the impact of SETD2 deficiency on protein synthesis, we pulse-labeled *K-Ctrl* and *K-Setd2^KO^* mice with the tRNA mimetic O-propargyl-puromycin (OP-Puro). OP-Puro is incorporated into actively translated proteins to allow for quantification of protein synthesis by flow cytometry or fluorescent microscopy and offers a method to quantify the rate of protein synthesis that is compatible with small sample sizes [12–14]. Combined staining for H3K36me3 and OP-Puro incorporation demonstrated a marked enhancement of overall protein synthesis in *K-Setd2^KO^* tumors (**Fig. 3A,B**). Additionally, while *KPY-Setd2^KO^* tumor cells were only slightly larger (3.8% higher FSC-A) and mRNA expression from the *Rosa26^LSL^-^YFP^* allele was similar to *KPY-Ctrl* tumor cells, *KPY-Setd2^KO^* tumor cells had an 18.5% increase in mean YFP fluorescence. This indicates a positive effect on YFP protein synthesis without affecting YFP mRNA production. (**Supplementary Fig. 4A,B**). These data suggest that SETD2 normally constrains the rate of protein synthesis in addition to the degree of OXPHOS. These functions, which would be expected to limit cellular proliferation, are consistent with the proliferation-driving effect of *Setd2* inactivation in KRAS-driven lung cancer.

**Figure 3:**
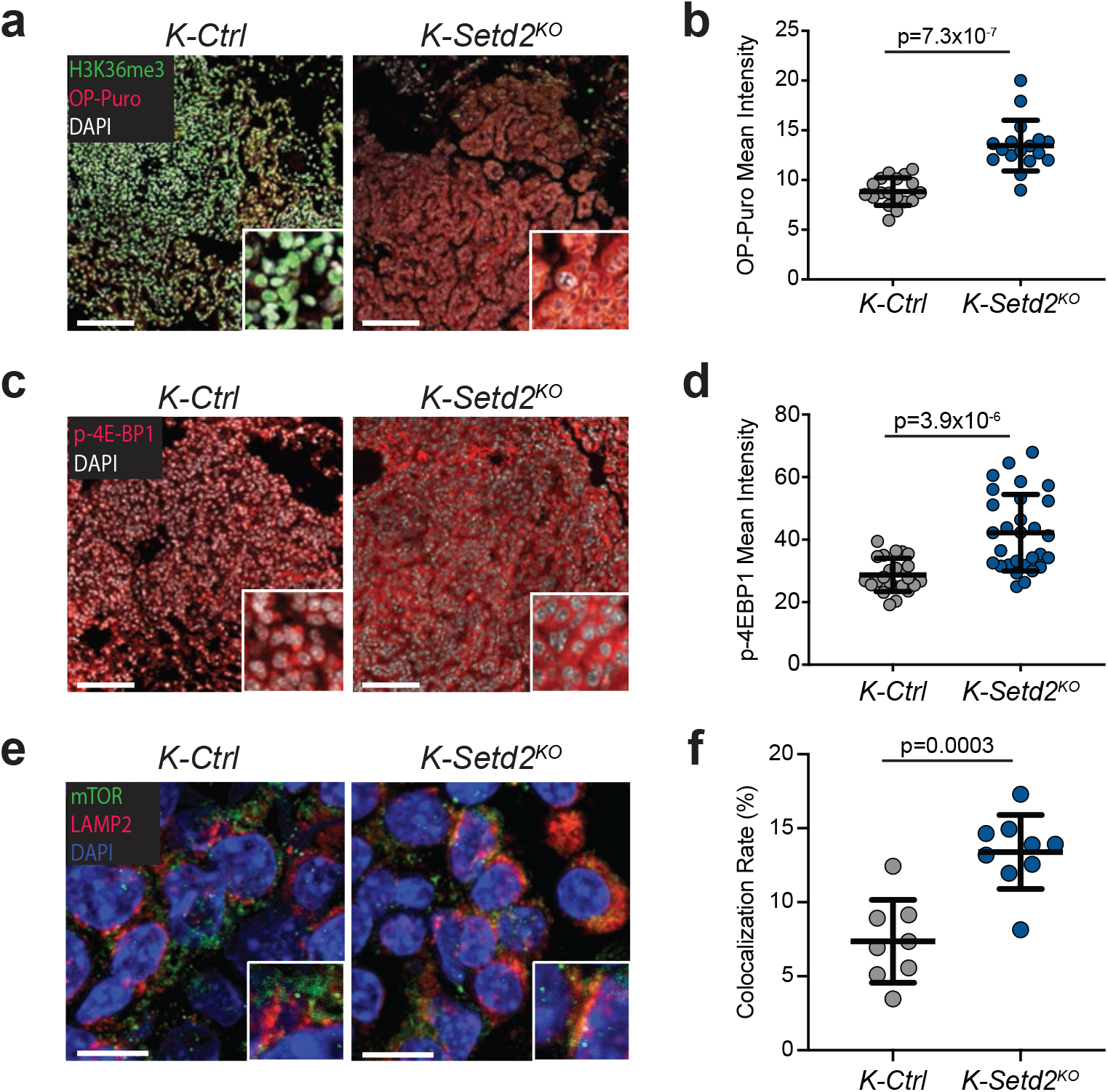
SETD2-deficient tumors have increased protein synthesis and high mTORC1 signaling. **a)** Representative images of o-propargyl-puromycin (OP-Puro) incorporation (red), H3K36me3 (green), and DAPI-stained nuclei (white) in *K-Ctrl* (left) and *KP-Setd2^KO^(right)* tumors. Scale bars=100 μm, insets are magnified 5x. **b)** Quantification of OP-Puro mean fluorescence intensity in *K-Ctrl* and *K-Setd2^KO^* tumors. Data represent the mean ± standard deviation. Data points represent individual tumors (*K-Ctrl*: n=18 tumors, n=3 mice, *K-Setd2^KO^*: n=17 tumors, n=3 mice). Significance determined by unpaired Student’s *t*-test. **c)** Representative images of phosphorylated 4E-BP1(T37/46) staining (red) in *K-Ctrl* (left) and *K*-*Setd2^KO^*(right) tumors. Nuclei are counterstained with DAPI (white). Scale bars=100 μm, insets are magnified 5x. **d)** Quantification of p-4E-BP1(T37/46) mean fluorescence intensity in *K-Ctrl* and *K-Setd2^KO^* tumors. Data represent the mean ± standard deviation. Data points represent individual tumors (*K-Ctrl*: n=24 tumors, n=3 mice, *K-Setd2^KO^*: n=29 tumors, n=3 mice). Significance determined by unpaired Student’s *t*-test. **e)** Representative images of co-immunofluorescence of mTOR (green) and LAMP2 (red) indicating localization of mTOR at the lysosome (yellow) in *K-Ctrl* (left) and *K-Setd2^KO^*(right) tumors. Nuclei are counterstained with DAPI (blue). Scale bars= 10 μm, inset is magnified 2X. **f)** Quantification of the percentage of colocalization between mTOR and LAMP2 in *K-Ctrl* and *K-Setd2^KO^* tumors. Data points represent individual tumors (*K-Ctrl*: n=8 tumors, n=3 mice, *K-Setd2^KO^*: n=9 tumors, n=3 mice). Significance determined by unpaired Student’s *t*-test.

Tightly linked with both protein synthesis and OXPHOS is the master nutrient sensing complex mTORC1 [15–17]. To determine whether SETD2 deficiency is associated with increased mTORC1 activity, we first evaluated human lung adenocarcinomas using reverse phase protein array (RPPA) data from the Cancer Proteome Atlas [18, 19]. An established marker of mTORC1 activity is the sequential phosphorylation of the translational repressor 4E-BP1, first at Thr37/Thr46 to prime subsequent phosphorylation at Thr70 and Thr65 [20, 21]. SETD2-deficient human tumors had significantly increased phosphorylation of 4E-BP1 at Thr70, while total 4E-BP1 levels were unchanged (**Supplementary Fig. 4C**). Additionally, there was a significant negative correlation between *SETD2* mRNA expression and phosphorylated 4E-BP1(T70) (**Supplementary Fig. 4D**). Consistent with these analyses of human datasets, we identified significantly increased mTORC1-dependent 4E-BP1(T37/46) phosphorylation in *K-Setd2^KO^* tumors compared to *K-Ctrl* tumors (**Fig. 3C,D**) [22]. Additionally, SETD2-deficient tumors had significantly higher levels of mTORC1 localized at the lysosome further indicating that SETD2 deficiency promotes mTORC1 activity (**Fig. 3E,F**).

Our discovery that mTORC1 signaling, protein synthesis, and mitochondrial OXPHOS are increased in SETD2-deficient tumors suggested that these processes may drive cell proliferation and thus offer therapeutic susceptibilities for *SETD2*-mutant cancers. To assess this possibility, we treated mice bearing established *K-Ctrl* and *K-Setd2^KO^* tumors daily for 4 weeks with either the mTORC1 inhibitor rapamycin, the mitochondrial complex I inhibitor IACS-10759, or the anti-diabetic biguanide phenformin which inhibits both complexes (**Fig. 4A**) [23–25]. All three therapeutics had little effect on *K-Ctrl* tumors. However, each treatment significantly suppressed the increased tumor growth of *K-Setd2^KO^* tumors (**Fig. 4B,C**). SETD2-deficient tumors were particularly sensitive to rapamycin treatment, which had a similar suppressive effect on tumor growth as phenformin which inhibits both mitochondrial complex I and mTORC1 signaling (**Fig. 4B,C**). Inhibition of mTORC1 and mitochondrial complex I activity resulted in significantly reduced cell proliferation, demonstrating that the proliferative impact of SETD2 inactivation is driven, at least in part, through these pathways (**Fig. 4D, Supplementary Fig. 5A**). Further, inhibition of mTORC1 and mitochondrial complex I, alone or in combination, did not result in significant levels of cell death indicating that cell death is not the cause of the decreased tumor growth observed (**Supplementary Fig. 5B**). Phenformin is a highly potent inhibitor of both mitochondrial complex I and mTORC1. This potency has led to significant toxicity and led to its clinical replacement for the treatment of type II diabetes with the related biguanide metformin, which is used widely and is epidemiologically associated with suppressing cancer incidence [26–28]. Therefore, we treated *K-Ctrl and K-Setd2^KO^* mice bearing established tumors with metformin for an extended period of 12 weeks, mimicking long-term metformin treatment. While metformin treatment had no impact on *K-Ctrl* tumor growth, metformin-treated *K-Setd2^KO^* tumors were significantly smaller and less proliferative than vehicle treated *K-Setd2^KO^* tumors (**Fig. 4E-G**).

**Figure 4:**
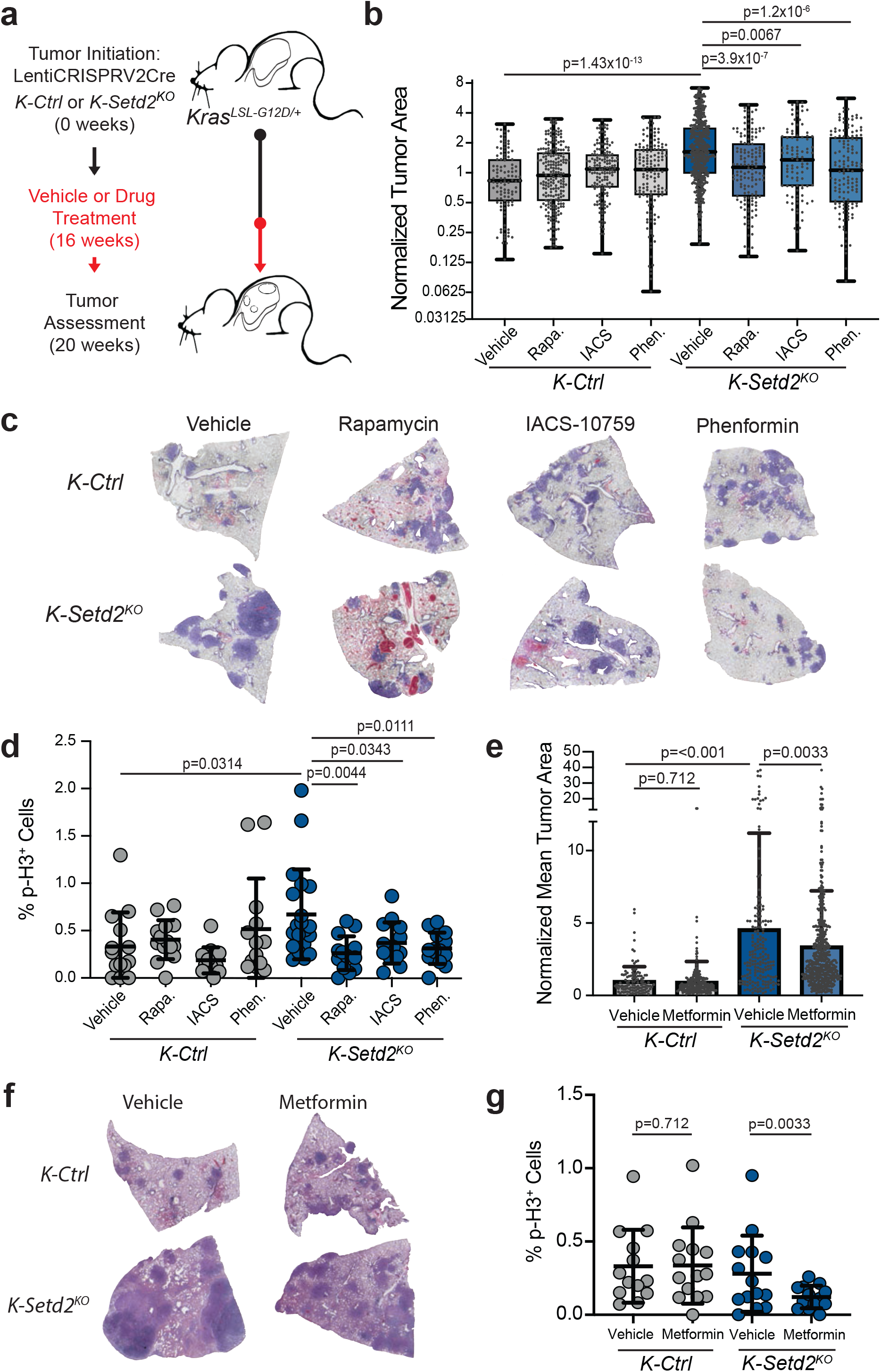
mTORC1 signaling and mitochondrial OXPHOS are required for the growth-promoting effects caused by SETD2-deficiency. **a)** Schematic of experiment whereby *K-Ctrl* or *Setd2^KO^* tumors were initiated using LentiCRISPRv2Cre. 16 weeks after tumor initiation (black line) mice were randomly assigned to vehicle or drug treatment regimen for 4 weeks (red line), at which point mice were sacrificed for assessment. For metformin treatment, mice were given the drug for 12 weeks prior to sacrifice. **b)** Quantification of mean tumor areas in vehicle-, rapamycin-, IACS-10759-, or phenformin-treated *K-Ctrl* or *K-Setd2^KO^* mice. Individual tumor sizes were normalized to the mean area of vehicle-treated control mice. Box and whisker plots indicate the median, lower quartile, upper quartile, maximum and minimum data points. Data points represent individual tumors (*K-Ctrl*: n=109 vehicle, n=216 rapamycin, n=155 IACS-10759, n=144 phenformin; *K-Setd2^KO^*: n=503 vehicle, n=142 rapamycin, n=99 IACS-10759, n=138 phenformin). Significance determined by unpaired Student’s *t*-test. **c)** Representative scans of tumor-bearing lobes from *K-Ctrl* or *K-Setd2^KO^* mice treated with vehicle, rapamycin, IACS-10759 or phenformin. **d)** Quantification of cell proliferation by the percentage of p-H3 positive cells in *K-Ctrl* or *K-Setd2^KO^* tumors treated with vehicle, rapamycin, IACS-10759 or phenformin. Data indicate the mean ± standard deviation. Data points represent individual tumors (*K-Ctrl*: n=14 vehicle, n=14 rapamycin, n=13 IACS-10759, n=13 phenformin; *K-Setd2^KO^*: n=20 vehicle, n=14 rapamycin, n=14 IACS-10759, n=14 phenformin). Significance determined by unpaired Student’s *t*-test. **e)** Quantification of mean tumor areas in vehicle- and metformin-treated *K-Ctrl* or *K-Setd2^KO^* mice. Individual tumor sizes normalized to the mean area of vehicle-treated control mice. Data represent the mean ± standard deviation. Data points represent individual tumors (*K-Ctrl*: n=115 vehicle, n=236 metformin; *K-Setd2^KO^*: n=193 vehicle, n=503 metformin). Significance determined by unpaired Student’s *t*-test. **f)** Representative scans of tumor-bearing lobes from *K-Ctrl* or *K-Setd2^KO^* mice treated with vehicle or metformin. **g)** Quantification of cell proliferation by the percentage of p-H3 positive cells in *K-Ctrl* and *K-Setd2^KO^* mice given vehicle treatment or metformin. Data indicate the mean ± standard deviation. Data points represent individual patient tumors (*K-Ctrl*: n=13 Vehicle, n=14 Metformin; *K-Setd2^KO^*: n=14 Vehicle, n=13 Metformin). Significance determined by unpaired Student’s *t*-test.

## Discussion

SETD2 is most commonly associated with its unique ability to catalyze histone H3 lysine 36 trimethylation (H3K36me3) along gene bodies during transcription [4]. However, multiple non-histone substrates have been identified which link SETD2 function with diverse roles in chromosome segregation (α-tubulin), interferon signaling (STAT1), or regulation of other chromatin modifying enzymes (EZH2) [5–7]. Additionally, H3K36me3 loss is implicated in the impairment of DNA repair processes, accurate splice site usage, transcriptional control, and DNA and RNA methylation [8–12]. Despite these numerous, characterized functions of SETD2, how SETD2 mediates lung tumor suppression remains unclear. Our study has identified a conserved enhancement of gene expression programs and functional markers associated with oxidative metabolism and protein synthesis in SETD2-deficient cancers. We demonstrate not only that signaling through each pathway is required for the proliferative benefit bestowed upon tumor cells following SETD2 inactivation, but that they also represent readily actionable therapeutic vulnerabilities for patients with SETD2-deficient tumors.

## Acknowledgements

We would like to thank H. Ehly and E. Morris for animal husbandry, B. Zuo for help with electron microscopy, A. Bedenbaugh for tissue processing, A. Durham and E. Radaelli for help with pathology, Bang-Jin Kim for assistance with confocal microscopy, and M. Winslow, K. Wellen, and S. Zhao for helpful discussions and critical reading of the manuscript.

## Methods

### Animal studies and treatment

Animal studies were performed under strict compliance with Institutional Animal Care and Use Committee at University of Pennsylvania (804774). *Kras^LSL-G12D^* mice (Jax stock number 008179), *Trp53^flox/flox^* mice (Jax stock number 008462), and *Rosa26^LSL-YFP/LSL-YFP^* mice have previously been described[23, 58, 59]. Mice are mixed B6J/ 129S4vJae. Mice were transduced with 6 × 10^4^ plaque forming units (PFUs) per mouse of LentiCRISPRv2Cre by endotracheal intubation as previously described[60]. LentiCRISPRv2Cre expressing sgRNAs targeting *GFP* or *β-Galactosidase* (BGal) were used as controls, while an sgRNA targeting *Setd2* was used for knockouts. For control sgRNAs, sgGFP was used to induce *K* tumors for RNA-sequencing, while sgBGal was used for all other experiments. The sgRNA sequences are: sgGFP – GGGCGAGGAGCTGTTCACCG, sgBGal – CACGTAGATACGTCTGCATC, and sgSetd2 – AATGGGCTGAGGTACGCCGT[14, 61]. Lentivirus production and titration was performed as described previously[14].

For drug experiments, OpenStandard Diet was formulated with Rapamycin (MedChem Express) at 15 mg kg^−1^ for a dose of 2 mg/kg/day or IACS-10759 (MedChem Express) at 37.5 mg kg^−1^ for a dose of 5 mg/kg/day by Research Diets. Mice were placed on treatment diet 4 weeks prior to analysis. For phenformin treatment, mice were given the drug by oral gavage daily on a 5 days on/2 days off schedule for 4 weeks. Mice were given 200 mg/ kg/day phenformin dissolved in water (Cayman Chemicals). For metformin treatment, metformin was placed in the drinking water of mice at 1.25 mg/ml for 12 weeks prior to sacrifice (Sigma-Aldrich). No statistical methods were used to predetermine sample sizes. The size of each animal cohort was determined by estimating biologically relevant effect sizes between control and treated groups and then using the minimum number of animals that could reveal statistical significance using the indicated tests of significance. All animal studies were randomized in ‘control’ or ‘treated’ groups, and roughly equal proportions of male and female animals were used. However, all animals housed within the same cage were generally placed within the same treatment group. For histopathological assessments of tumor size, researchers were blinded to sample identity and group.

### Immunohistochemistry and immunofluorescence

Lung and tumor tissues were dissected into 10% neutral-buffered formalin overnight at room temperature before dehydration in a graded alcohol series. Paraffin-embedded and H&E-stained histological sections were produced by the Penn Molecular Pathology and Imaging Core. Immunostaining for H3K36me3 (Abcam, ab9050, 1:1000), p-4E-BP1(T37/46) (Cell Signaling Technology, cs2855, 1:100), mTOR (Cell Signaling Technology, cs2983, 1:100), LAMP2 (Abcam, ab13524, 1:100) and p-H3 (Cell Signaling Technology, cs9701, 1:500) were performed after citrate-based antigen retrieval. H3K36me3 alone was assessed by immunohistochemistry using ABC reagent (Vector Laboratories, PK-4001) and ImmPACT DAB (Vector Laboratories, SK-4105) according to the product instructions. P-4E-BP1(T37/46) was assessed by immunofluorescence using a biotinylated secondary antibody (Vector Laboratories, PK-4001) according to product instructions, and Streptavidin-conjugated Alexa594 (Thermo Fisher S11227, 1:200). Colocalization of mTOR and LAMP2 was determined using an anti-Rat Alexa647 antibody (Thermo Fisher A21247, 1:200) to detect LAMP2, and a biotinylated anti-Rabbit secondary antibody (Vector Laboratories, PK-4001) followed by Streptavidin-conjugated Alexa488 (Thermo Fisher S32354, 1:200) to detect mTOR.

Immunohistochemistry and immunofluorescence were both performed on paraffin-embedded sections following the same antigen-retrieval protocol. Sections were incubated in primary antibody overnight at 4 °C, secondary antibody for 1 hour at room temperature, and for immunofluorescence Streptavidin-conjugated fluorophore for 1 hour at room temperature in the dark.

For TUNEL staining, tissues were deparaffinized and then permeabilized with 0.1% sodium citrate and 0.1% Triton-X in PBS for 8 minutes. FITC-conjugated TUNEL labeling mix (Millipore Sigma, 11684795910) was added to permeabilized tissue sections and incubated for 1 hour at 37 °C in the dark. For all immunofluorescence staining, nuclei were stained using 5 mg/ ml DAPI at a 1:1000 dilution for 10 minutes, and then slides were mounted with Fluoro-Gel (EMS, 17985-50).

### O-Propargyl-Puromycin Analysis

For the quantification of protein translation, mice were injected intraperitoneally (IP) with 200 μl of a 10 mM solution of OP-Puro dissolved in PBS as previously described[24]. 1 hour after OP-Puro IP injection, mice were sacrificed and lungs were formalin fixed and paraffin embedded as described above. Co-immunofluorescence for H3K36me3 and OP-Puro was performed to quantify translation rates in tumors lacking SETD2 activity. Antigen retrieval was performed using a solution of 20 μg/ml proteinase K in TE Buffer (pH 8) at 37 °C for 10 minutes. A click chemistry reaction was then performed for 30 minutes at room temperature in the dark to conjugate Alexa594 to incorporated OP-Puro according to product instructions (Thermo Fisher, C10429, beginning at Step 5.1). Samples were kept in the dark for all further steps. Samples were treated with avidin and biotin blocking steps for 20 minutes each (Vector Laboratories, SP-2001), and a 30-minute protein block (Dako, X090930-2) before incubating with H3K36me3 primary antibody (Abcam, ab9050) at 1:300 overnight at 4 °C. H3K36me3 was then detected using a biotinylated secondary antibody for 1 hour (Vector Laboratories, PK-4001) followed by streptavidin-conjugated Alexa488 for 1 hour at a 1:200 dilution (Thermo Fisher, S-32354). Nuclei were stained using 5 mg/ml DAPI at a 1:1000 dilution for 10 minutes, and then slides were mounted with Fluoro-Gel (EMS, 17985-50).

### Histological quantification

The analysis of mTOR/LAMP2 colocalization was performed using a Leica TCS SP5 II confocal microscope. Z-stack projections of confocal images taken of control and SETD2-deficient tumors were analyzed. For the quantification of PLA individual loci were counted from z-stack projections. The LAS X colocalization tool was used for the quantification of mTOR/ LAMP2 colocalization, All other photomicrographs were captured on a Leica DM16000B inverted light and fluorescence microscope, and ImageJ software was used for subsequent histological quantifications[64]. For the quantification of staining intensity for OP-Puro and p-4EBP1(T37/46), single fluorescent channel images were obtained and the mean fluorescence intensity of staining for each tumor was quantified in ImageJ. Great care was made to ensure that background signal from blood vessels, or empty spaces were excluded from the analysis. For the quantification of tumor sizes under varying conditions including drug treatments, a tile scan of each mouse lung was obtained using the Leica DM1600B microscope and tumor area was measured in ImageJ. Tumor areas were then normalized to the mean area of a sgCtrl, vehicle treatment tumor. For the quantification of p-H3 staining, total nuclei in each tumor were counted using the IHC Profiler plugin for ImageJ, and p-H3-expressing nuclei were counted in ImageJ using the Cell Counter plugin[65]. For all histological analyses each data point represents an individual tumor.

### Flow Cytometry

Tumors were microdissected directly from the lungs of *Kras^LSL-G12D/+^;Trp53^flox/flox^*; *Rosa26^LSL-YFP/LSL-YFP^(KPY)* mice and individually placed in 500 μl of tumor digestion buffer consisting of PBS containing 10 mM HEPES pH 7.4, 150 mM NaCl, 5 mM KCl, 1 mM MgCl_2_, and 1.8 mM CaCl_2_, along with freshly added Collagenase 4 (Worthington 100 mg/ml solution, 20 μl per ml of digestion buffer) and DNase I (Roche 10 mg/ml solution, 4 μl per ml of digestion buffer). Tumors were manually disassociated using scissors, and then placed in a 4 °C shaker for 1 hour at 250 rpm. Digested tumors were then filtered into strainer-cap flow tubes (Corning, 352235) containing 1 ml of horse serum (Thermo Fisher, 16050122) to quench the digestion reaction. Cells were spun down at 200 g for 5 minutes with the cap in place to obtain all cells. The supernatant was aspirated, cells were washed once with PBS and then resuspended with a given mitochondrial dye to stain for 30 minutes at 37 °C. To quantify mitochondrial volume, cells were incubated with 50 nM MitoTracker Deep Red FM (Thermo Fisher, M22426), combined with 50 μM of CCCP (Thermo Fisher, M34151) to eliminate confounding effects of mitochondrial membrane potential differences. To quantify mitochondrial membrane potential cells were incubated with 20 nM MitoProbe DiIC1(5) (Thermo Fisher, M34151). To quantify mitochondrial oxidative potential cells were incubated with 100 nM MitoTracker Red CM-H2XRos (Thermo Fisher, M7513). To quantify mitochondrial ROS cells were incubated with 5 μM MitoSOX Red (Thermo Fisher, M36008). After 30 minutes of staining, cells were washed twice with PBS and then resuspended in 100 μl of staining solution of FACS buffer containing biotinylated antibodies against CD31 (BD Biosciences, 558737, 1:100), CD45 (BD Biosciences, 553078, 1:200), and Ter-119 (BD Biosciences, 553672, 1:100), for 25 minutes at a 4 °C. Cells were washed twice and resuspended in 100 μl of streptavidin-conjugated APC-eFluor 780 (Thermo Fisher, 47-4317-82) for 20 minutes. Finally, cells were washed twice and resuspended in FACS buffer containing DAPI at a 1:1000 dilution. Flow cytometry was performed using an Attune NxT flow cytometer (Thermo Fisher), and gating was performed to exclude doublets, dead cells, YFP-cells and non-epithelial contaminating cell types (see **Supplementary Fig. 2c**). Mitochondrial properties were then quantified by measuring the median fluorescence intensity of a given dye in live, YFP+ tumor cells. For the quantification of YFP protein expression the mean fluorescence intensity of YFP was quantified for each sample by flow cytometry. For all flow cytometry analyses each data point represents an individual tumor.

### RNA sequencing and human dataset analysis

For gene expression analysis in human tumors, RNA-sequencing data was obtained from the Cancer Genome Atlas lung adenocarcinoma dataset[2]. Due to the relative infrequency of *SETD2* mutations, a comparison of mutant and wildtype cases had insufficient statistical power to draw meaningful conclusions. Therefore, mRNA expression data was extracted by the Penn Institute for Biomedical Informatics, and a pearson correlation score was calculated comparing the expression of *SETD2* to all other genes. All genes were ranked in order according to the genes most negatively correlated with *SETD2* expression to most positively correlated, and this rank list was used to perform Gene Set Enrichment Analysis (GSEA) examining GO biological processes using the molecular signature database (MSigDB)[66, 67]. For gene expression analysis in *K-Ctrl* and *K-Setd2^KO^* tumors, analysis was performed on previous sequencing results[14]. For gene expression analysis in *KPY-Ctrl* and *KPY-Setd2^KO^* tumors, tumors were microdissected away from normal lung tissue, and digested into a single cell suspension as described above. Live, YFP+ tumor cells were isolated by cell sorting, spun down and flash frozen in liquid nitrogen. RNA was extracted using the RNeasy Plus Micro kit (Qiagen, Catalog #74034) using 350 μl of RLT Plus and the QIAshredder columns as per manufacturer’s instructions. Total RNA quantity was measured using the Qubit RNA HS assay kit (ThermoFisher, Catalog #Q32852) and RNA quality was measured using a BioAnalyzer RNA 6000 Nano assay (Agilent, Catalog #5067-1511). Sequencing libraries were prepared on the Illumina NeoPrep and subjected to 75-bp single-end sequencing on the Illumina NextSeq 500 platform. Fastq files for each sample were aligned against the mouse genome, build GRCm38.p5, using Salmon (v0.8.2)[68].

Differentially expressed genes were identified with DESeq2 (v1.17.0) and ranked according to the Stat value which considers both the significance, fold-change and directionality of the gene expression change[69]. This rank of genes most upregulated upon *Setd2* loss was then used to perform GSEA examining GO biological processes using the MSigDB. GSEA network plots of the top 20 pathways negatively correlated with *SETD2* expression in both human and mice were then generated as previously described[19]. A graphical depiction of the network plot was then generated using Gephi v.0.9.2, and gene sets were characterized according to their functions[70]. To quantify YFP RNA expression the total YFP read count was quantified from RNA-sequencing data of *KPY* tumors using Salmon (v0.8.2).

To analyze 4E-BP1 protein levels and phosphorylation in human tumors, level 4 RPPA data from human lung adenocarcinomas was extracted from the Cancer Proteome Atlas website[30, 31]. RPPA z-scores were matched with gene expression and genetic information from each sample represented in the TCGA lung adenocarcinoma dataset[2]. The RPPA z-score for phosphorylated 4E-BP1(T70) was compared to *SETD2* mRNA expression z-scores for each sample. The levels of total 4E-BP1 and phosphorylated 4E-BP1(T70) were also compared between tumor samples with wildtype *SETD2*, and *SETD2* deficiency (defined as tumors containing either an inactivating mutation in *SETD2*, homozygous loss of the gene, or a loss of 1 copy of *SETD2* along with a mRNA z-score <-0.5).

### Electron Microscopy

Tumors were microdissected directly from the lungs of mice and then the tissue was divided in half. One portion of each tumor was fixed for IHC to determine the H3K36me3-status of the given tumor as described above, while the other portion was fixed overnight in an osmium solution obtained from the Penn Electron Microscopy Resource Lab (EMRL), and then submitted to the EMRL for further tissue processing and staining with uranyl acetate and lead citrate. Transmission electron microscopy (TEM) was then performed using a JEOL JEM-1010 for both control and SETD2-deficient tumor samples. Images were taken at 60,000 to 150,000 X magnification, and mitochondrial properties were then quantified using ImageJ, normalizing to the magnification of the image. Mitochondrial size was in Image J, while mitochondrial number was quantified by counting the number of mitochondria per field of view across multiple 15,000 X magnification images. Mitochondrial cristae width was quantified in Image J by drawing a perpendicular line between the inner membranes of cristae, and then quantifying the resulting distance. Mitochondrial electron density was quantified by measuring the mean pixel darkness of the mitochondrial matrix in ImageJ and normalizing this to the mean pixel darkness of the surrounding cytoplasm. Mitochondrial cristae density was quantified by counting the number of cristae in an individual mitochondrion and then dividing by the mitochondrial area.

### Seahorse XF Cell Mito Stress Analysis

Oxidative respiration was measured using XF Cell Mito Stress Test Kit (Agilent Technologies, 103015-100). 1 × 10^4^ cells per well were seeded on an XF96 Cell Culture Microplate. Microplate was incubated for 24h at 37C. Seahorse XF96 FluxPak sensor cartridge was hydrated with 200 μl of Seahorse Calibrant in a non-CO_2_ incubator at 37C overnight. After 24h, cells were incubated with base medium (Agilent Technologies, 102353-100) containing 2 mM L-glutamine, 1 mM sodium pyruvate, and 10 mM glucose in a non-CO_2_ incubator at 37C for 45 min prior to assay. Oxygen consumption rate (OCR) was measured by XFe96 extracellular flux analyzer with sequential injections of 1 μM oligomycin, 1 μM FCCP, and 0.5 μM rotenone/antimycin A. After the run, cells were lysed with 15 μl RIPA buffer and protein concentration was quantified using Pierce BCA Protein Assay Kit (Thermo Fisher Scientific, 23225). OCR measurements were normalized to the protein concentration in each well.

### Statistical analysis

All analyses were performed using Graphpad Prism (v.8.1.1). For all analyses of mitochondrial properties, immunofluorescence, percentage of p-H3^+^ cells, normalized tumor areas and RPPA analysis comparing *SETD2* wildtype and deficient tumors, unpaired Student’s *t*-tests were performed. For the comparison of *SETD2* mRNA expression and p-4E-BP1(T70) levels by RPPA, a linear regression analysis was performed. Outliers were excluded from rapamycin, IACS-10759 and phenformin experiments using the ROUT method with a Q of 0.1%.

---

**Supplementary Fig. 1:**
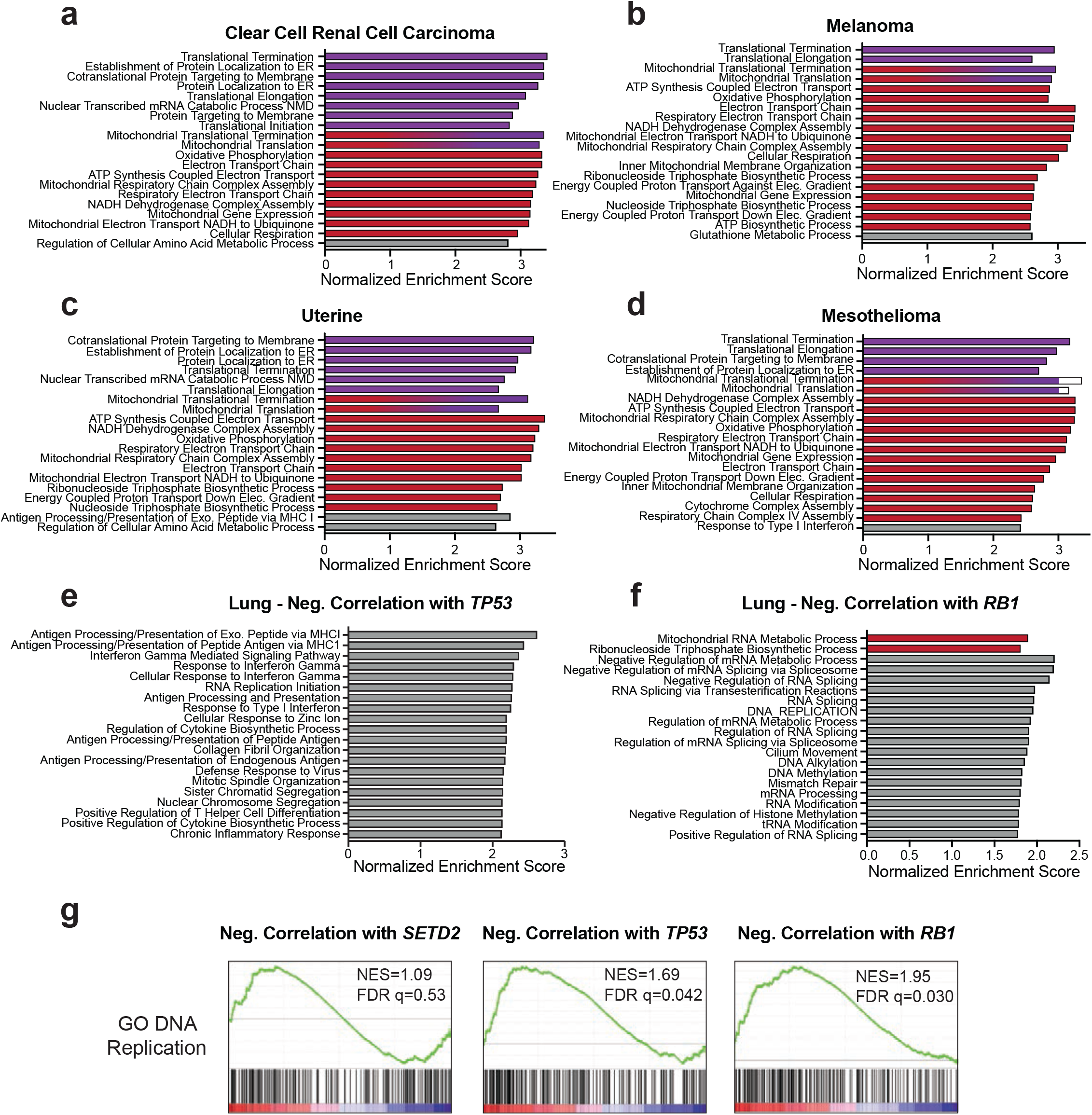
SETD2 deficiency promotes protein synthesis and OXPHOS gene expression programs in multiple human cancer types. Gene sets are categorized as mitochondrial (blue), ribosomal (green) or other (gray) according to the genes represented. Gene sets containing both mitochondrial and ribosomal genes are blue with green diagonal lines. The size of the bar indicates the normalized enrichment score (NES) of the gene set. **a)** The top 20 gene sets in human clear cell renal cell carcinoma correlated with low expression of *SETD2*. **b)** The top 20 gene sets in human melanoma correlated with low expression of *SETD2*. **c)** The top 20 gene sets in human uterine cancer correlated with low expression of *SETD2*. **d)** The top 20 gene sets in human mesothelioma correlated with low expression of *SETD2*. **e)** The top 20 gene sets in human lung adenocarcinoma correlated with low expression of *TP53*. **f)** The top 20 gene sets in human lung adenocarcinoma correlated with low expression of *RB1*. **g)** Gene set enrichment analysis plots of the GO DNA Replication gene negatively correlated with *SETD2, TP53* or *RB1* gene expression in human lung adenocarcinomas.

**Supplementary Fig. 2:**
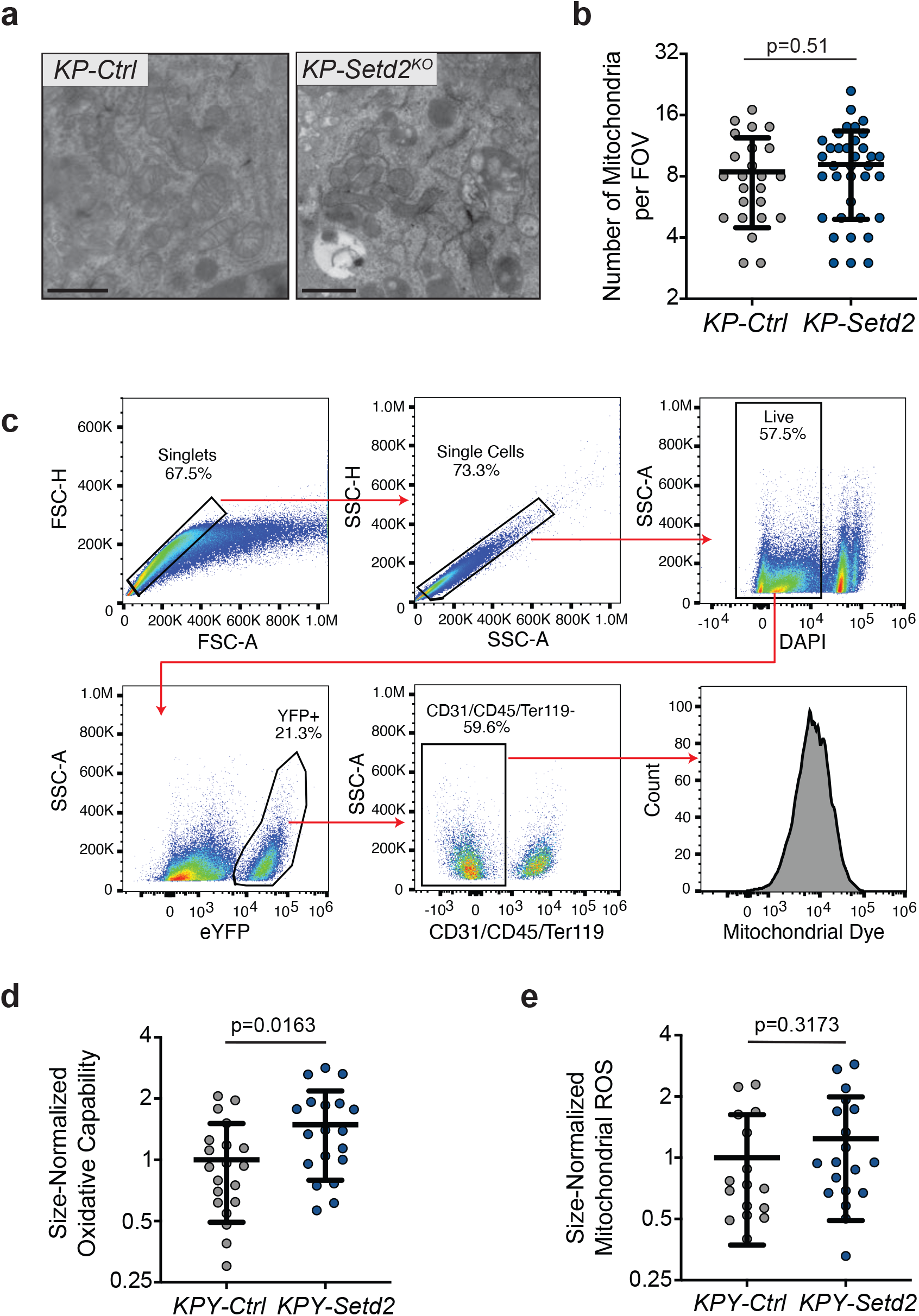
Characterization of mitochondrial properties in *KP-Ctrl* and *KP-Setd2^KO^* tumors. **a)** Representative EM photomicrographs demonstrating that *KP-Setd2^KO^* tumors have similar numbers of mitochondria, that are smaller and more electron dense, than *KP-Ctrl* tumors. Scale bars =2 μm. **b)** Quantification of the number of mitochondria per field of view from electron micrographs of 15,000X magnification. Data indicate the mean ± standard deviation. Data points represent individual tumors (*KP-Ctrl*: n=24 micrographs, n=3 mice, *KP-Setd2^KO^*: n=36 micrographs, n=3 mice). Significance determined by unpaired Student’s *t*-test. **c)** Gating strategy for dye-based, *ex vivo* mitochondrial analyses in isolated tumor cells from *KPY* mice. Doublets are excluded based on FSC and SSC properties. DAPI-positive dead cells and YFP-negative cells are excluded. CD31/CD45/Ter-119-positive non-epithelial cell types are excluded. Remaining YFP-positive tumor cells are analyzed for mitochondrial dye fluorescence intensity. **d)** Quantification of the oxidative capability of mitochondria within *KPY-Ctrl* and *KPY-Setd2^KO^* tumor cells by the median fluorescence intensity of MitoTracker Red CM-H2-XRos staining by flow cytometry and normalized to the mitochondrial mass of each sample. Data represent the mean ± standard deviation. Data points represent individual tumors (*KPY-Ctrl*: n=20 tumors, n=2 mice, *KPY-Setd2^KO^*: n=19 tumors, n=2 mice). Significance determined by unpaired Student’s *t*-test. **e)** Quantification of mitochondrial reactive oxygen species (ROS)/superoxide levels within *KPY-Ctrl* and *KPY-Setd2^KO^* tumor cells by the median fluorescence intensity of MitoSOX Red staining by flow cytometry and normalized to the mitochondrial mass of each sample. Data represent the mean ± standard deviation. Data points represent individual tumors (*KPY-Ctrl*: n=16 tumors, n=2 mice, *KPY-Setd2^KO^*: n=19 tumors, n=2 mice). Significance as determined by unpaired Student’s *t*-test.

**Supplementary Fig. 3.**
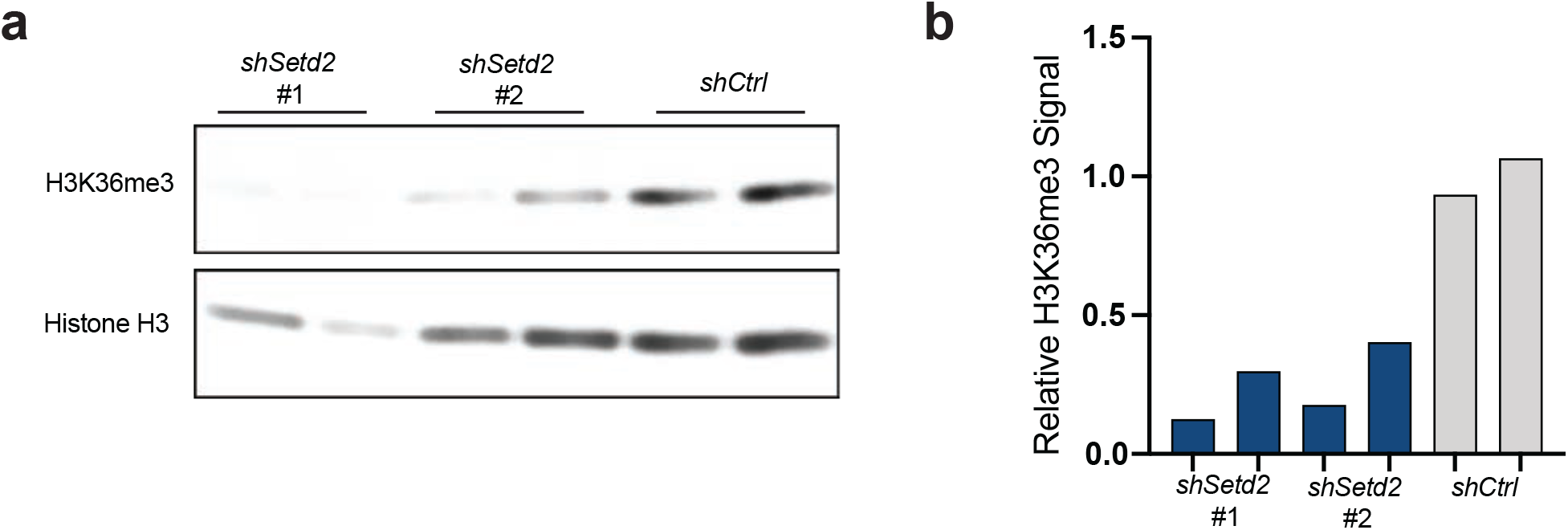
Generation of SETD2 shRNA human cell lines. **a)** Immunoblot analysis on histones derived from H2009 sh*Setd2* and sh*Ctrl* cell lines for H3K36me3. Histone H3 is used as a loading control. **b)** Quantification of immunoblot from (a). H3K36me3 signal is normalized to total H3 for each lane.

**Supplementary Fig. 4:**
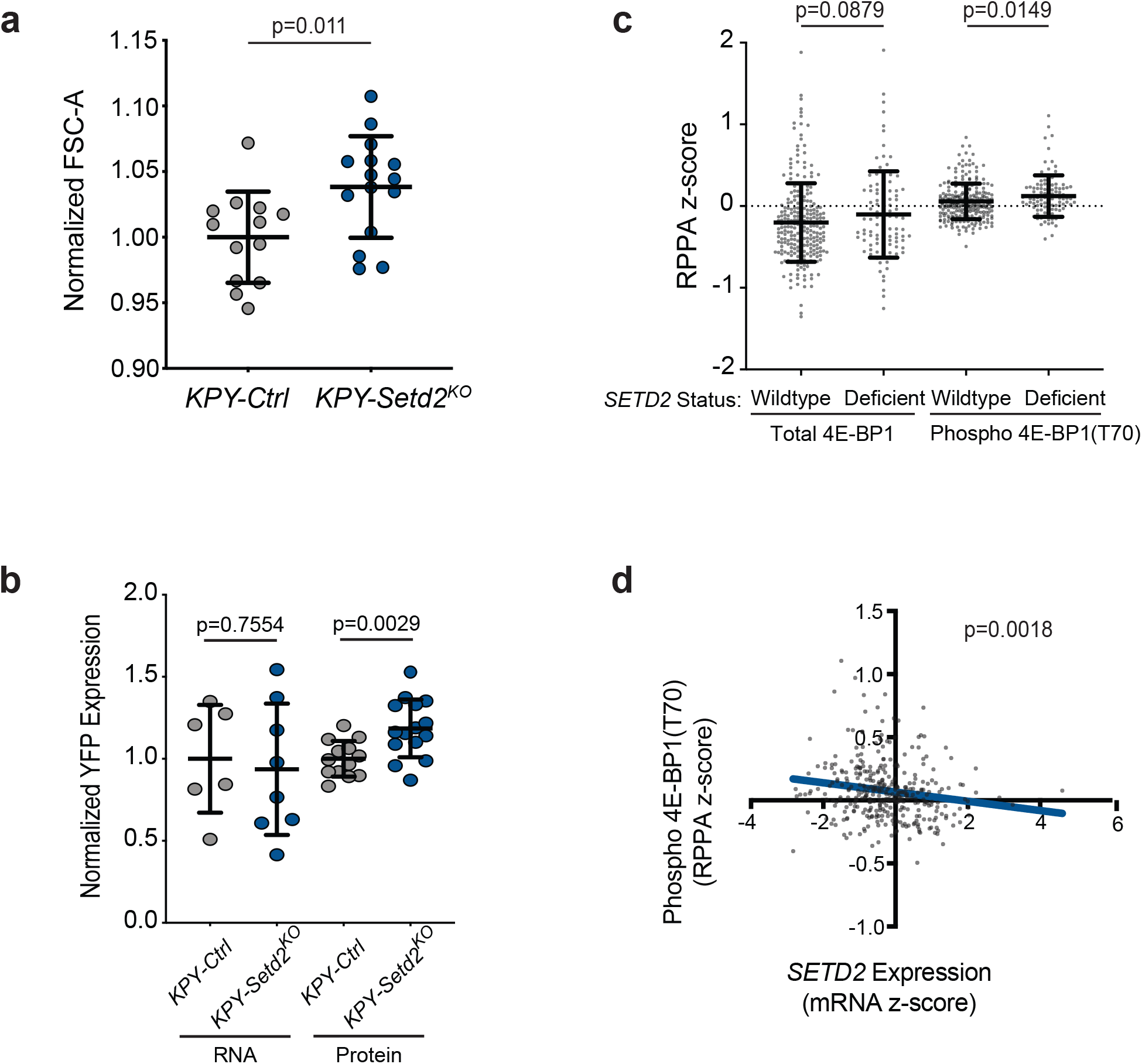
mTORC1 signaling and protein synthesis are heightened in SETD2-deficient tumors. **a)** Quantification of mean cell size in *KPY-Ctrl* and *KPY-Setd2^KO^tumors* by forward scatter area (FSC-A) by flow cytometry. Data indicate the mean ± standard deviation. Data points represent individual tumors (*KPY-Ctrl*: n=13 tumors, n=7 mice, *KPY-Setd2^KO^*: n=15 tumors, n=8 mice). Significance determined by unpaired Student’s *t*-test. **b)** Normalized expression of YFP RNA and protein in *KPY-Ctrl* and *KPY-Setd2^KO^* tumors. Data indicate the mean ± standard deviation. Data points represent individual tumors (RNA, *KPY-Ctrl*: n=6 tumors, n=4 mice, *KPY-Setd2^KO^*: n=8 tumors, n=5 mice. Protein, *KPY-Ctrl*: n=13 tumors, n=7 mice, *KPY-Setd2^KO^*: n=15 tumors, n=8 mice). Significance determined by unpaired Student’s *t*-test. **c)** Quantification of total 4E-BP1 and phosphorylated 4E-BP1(T70) by RPPA in human lung adenocarcinomas with wildtype *SETD2* or *SETD2* deficiency. Data indicate the mean ± standard deviation. Data points represent individual patient tumors (wildtype *SETD2*: n=253 tumors, *SETD2* deficiency: n=103 tumors). Significance determined by unpaired Student’s *t*-test. **d)** Analysis of the relationship between *SETD2* gene expression (mRNA z-score) and phosphorylated 4E-BP1(T70) as quantified by RPPA in human lung adenocarcinomas. Data points represent individual patient tumors (n=357 tumors). Significance determined by linear regression analysis and the line of best fit is shown.

**Supplementary Fig. 5:**
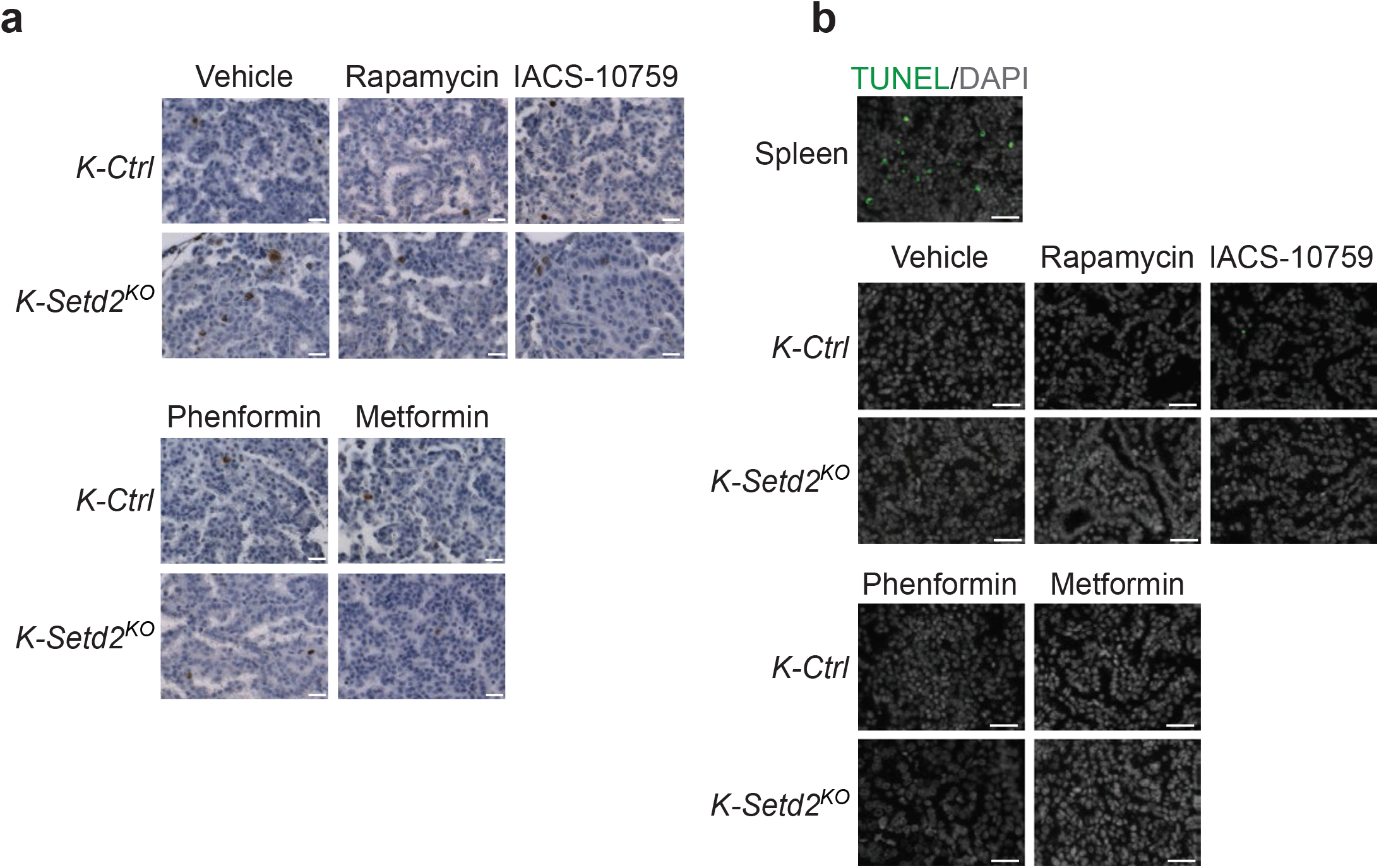
Inhibiting mTORC1 signaling or mitochondrial activity reduces cell proliferation in SETD2-deficient tumors. **a)** Representative images from p-H3 staining of *K-Ctrl* and *K-Setd2^KO^* tumors from mice treated with vehicle, rapamycin, IACS-10759, phenformin or metformin. Scale bars= 25 μm. **b)** Representative images from TUNEL staining of *K-Ctrl* and *K-Setd2^KO^* tumors from mice treated with vehicle, rapamycin, IACS-10759, phenformin or metformin. Positive control for TUNEL is a murine spleen. TUNEL is marked by green, DAPI-stained nuclei are in white. Scale bars= 300 μm.

## Bibliography

1. Kandoth, C., et al., Mutational landscape and significance across 12 major cancer types. Nature, 2013. 502(7471): p. 333–339.

2. Comprehensive molecular profiling of lung adenocarcinoma. Nature, 2014. 511(7511): p. 543–50.

3. Zehir, A., et al., Mutational landscape of metastatic cancer revealed from prospective clinical sequencing of 10,000 patients. Nature medicine, 2017. 23(6): p. 703–713.

4. Rogers, Z.N., et al., A quantitative and multiplexed approach to uncover the fitness landscape of tumor suppression in vivo. Nat Methods, 2017. 14(7): p. 737–742.

5. Walter, D.M., et al., Systematic In Vivo Inactivation of Chromatin-Regulating Enzymes Identifies Setd2 as a Potent Tumor Suppressor in Lung Adenocarcinoma. Cancer Res, 2017. 77(7): p. 1719–1729.

6. Rogers, Z.N., et al., Mapping the in vivo fitness landscape of lung adenocarcinoma tumor suppression in mice. Nat Genet, 2018. 50(4): p. 483–486.

7. Rasool, R.U., et al., CDK7 inhibition suppresses Castration-Resistant Prostate Cancer through MED1 inactivation. Cancer Discovery, 2019: p. CD-19-0189.

8. Cogliati, S., et al., Mitochondrial cristae shape determines respiratory chain supercomplexes assembly and respiratory efficiency. Cell, 2013. 155(1): p. 160–171.

9. Cogliati, S., J.A. Enriquez, and L. Scorrano, Mitochondrial Cristae: Where Beauty Meets Functionality. Trends in Biochemical Sciences, 2016. 41(3): p. 261–273.

10. Nielsen, J., et al., Plasticity in mitochondrial cristae density allows metabolic capacity modulation in human skeletal muscle. The Journal of Physiology, 2017. 595(9): p. 2839–2847.

11. Srinivas, S., et al., Cre reporter strains produced by targeted insertion of EYFP and ECFP into the ROSA26 locus. BMC Dev Biol, 2001. 1: p. 4.

12. Liu, J., et al., Imaging protein synthesis in cells and tissues with an alkyne analog of puromycin. Proceedings of the National Academy of Sciences of the United States of America, 2012. 109(2): p. 413–418.

13. Signer, R.A.J., et al., Haematopoietic stem cells require a highly regulated protein synthesis rate. Nature, 2014. 509(7498): p. 49–54.

14. Iwasaki, S. and N.T. Ingolia, The Growing Toolbox for Protein Synthesis Studies. Trends in biochemical sciences, 2017. 42(8): p. 612–624.

15. Cunningham, J.T., et al., mTOR controls mitochondrial oxidative function through a YY1-PGC-1alpha transcriptional complex. Nature, 2007. 450(7170): p. 736–40.

16. Saxton, R.A. and D.M. Sabatini, mTOR Signaling in Growth, Metabolism, and Disease. Cell, 2017. 168(6): p. 960–976.

17. de la Cruz López, K.G., et al., mTORC1 as a Regulator of Mitochondrial Functions and a Therapeutic Target in Cancer. Frontiers in Oncology, 2019. 9(1373).

18. Li, J., et al., TCPA: a resource for cancer functional proteomics data. Nat Methods, 2013. 10(11): p. 1046–7.

19. Li, J., et al., Explore, Visualize, and Analyze Functional Cancer Proteomic Data Using the Cancer Proteome Atlas. Cancer Res, 2017. 77(21): p. e51–e54.

20. Gingras, A.C., et al., Hierarchical phosphorylation of the translation inhibitor 4E-BP1. Genes Dev, 2001. 15(21): p. 2852–64.

21. Qin, X., B. Jiang, and Y. Zhang, 4E-BP1, a multifactor regulated multifunctional protein. Cell cycle (Georgetown, Tex.), 2016. 15(6): p. 781–786.

22. Dowling, R.J.O., et al., mTORC1-mediated cell proliferation, but not cell growth, controlled by the 4E-BPs. Science (New York, N.Y.), 2010. 328(5982): p. 1172–1176.

23. Shackelford, D.B., et al., LKB1 inactivation dictates therapeutic response of non-small cell lung cancer to the metabolism drug phenformin. Cancer Cell, 2013. 23(2): p. 143–58.

24. Xie, J., X. Wang, and C.G. Proud, mTOR inhibitors in cancer therapy. F1000Research, 2016. 5: p. F1000 Faculty Rev-2078.

25. Molina, J.R., et al., An inhibitor of oxidative phosphorylation exploits cancer vulnerability. Nat Med, 2018. 24(7): p. 1036–1046.

26. Misbin, R.I., The Phantom of Lactic Acidosis due to Metformin in Patients With Diabetes. Diabetes Care, 2004. 27(7): p. 1791.

27. Lee, M.-S., et al., Type 2 diabetes increases and metformin reduces total, colorectal, liver and pancreatic cancer incidences in Taiwanese: a representative population prospective cohort study of 800,000 individuals. BMC Cancer, 2011. 11(1): p. 20.

28. Currie, C.J., et al., Mortality after incident cancer in people with and without type 2 diabetes: impact of metformin on survival. Diabetes Care, 2012. 35(2): p. 299–304.

29. Mootha, V.K., et al., PGC-1alpha-responsive genes involved in oxidative phosphorylation are coordinately downregulated in human diabetes. Nat Genet, 2003. 34(3): p. 267–73.

